# Single-Molecule Dwell Times in Biomolecular Condensates

**DOI:** 10.64898/2026.06.29.735418

**Authors:** Fengshuo Yang, Roumita Moulick, Cailing Wang, Margaret L. Rodgers, Sarah A. Woodson, Yaojun Zhang

## Abstract

Biomolecular condensates are dynamic, membrane-free compartments that continuously exchange molecules with their surroundings. The dwell time, defined as the time a molecule remains inside a condensate between entry and exit, determines how extensively the molecule can explore the dense phase and encounter potential binding partners or reaction sites, thereby modulating condensate function. Motivated by our single-molecule measurements of RNA dwell times, we developed an analytical theory to understand dwell-time distributions in biomolecular condensates. Our theory predicts that the dwell-time distributions generally exhibit an early-time power-law regime followed by a late-time exponential tail. The form of the distribution encodes the rate-limiting mechanism of molecular escape: dense-phase diffusion-limited transport feature a −1.5 power law with an exponential tail set by a diffusion timescale, whereas interfacial barrier-crossing-limited transport feature a − 0.5 power law with a decay governed by a barrier-crossing timescale. These distinct signatures provide a direct readout of the physical processes that control molecular retention in condensates, with implications for both natural and synthetic condensates.

## I. INTRODUCTION

Biomolecular condensates have emerged as a central paradigm for how living cells use collective physical behavior to organize matter [1, 2]. This mode of organization underlies a wide range of cellular processes, from gene regulation to signaling, metabolism, and stress responses [3–6], while its dysregulation is implicated in disease [7, 8]. The nature of phase separation makes condensates intrinsically open and dynamic: they continuously exchange components with the dilute phase and assemble or dissolve quickly in response to cellular cues [9–11]. By tuning condensate dynamics, cells can regulate biochemical reactions, molecular accessibility, and the spatiotemporal organization of condensates [12–15]. Understanding condensate dynamics is therefore central to understanding their function.

The dynamical properties of condensates have been probed using a broad set of experimental approaches, including fluorescence recovery after photobleaching (FRAP) [10], fluorescence correlation spectroscopy (FCS) [16, 17], and passive and active microrheology [18].

Among these, single-molecule tracking (SMT) is particularly informative: by following individual fluorescently labeled molecules, SMT can resolve heterogeneity and rare events that are obscured in ensemble measurements, providing microscopic insights into molecular transport and interactions in condensates both *in vitro* and in cells [19–21].

A key SMT observable is the dwell time, defined as the time a molecule resides inside a condensate between entry and exit events (Fig. 1). Dwell times are biologically meaningful because they quantify how long molecules are retained in condensates for reaction, processing, or assembly. For example, in an engineered condensate model, the mean dwell time of the intrinsically disordered region (IDR) of protein TAF15 has been reported to be much shorter than that of oligomerized TAF15 IDR (10 s versus 64 s) [22], illustrating how oligomerization can serve as a regulatory knob to tune molecular retention within condensates. Another example comes from DNA double-strand break repair: SMT in live yeast cells estimated the mean residence time of the recombination mediator Rad52 in repair foci to be around 240 ms [23]. This rapid turnover could help prevent prolonged sequestration and thereby allow Rad52 to quickly redistribute among concurrent DNA damage sites. In the context of ribosome biogenesis, the mean residence time of rRNA transcripts in nucleoli is about 45 min [24]. Such a long dwell time is thought to ensure proper folding and processing of nascent rRNAs by enzymes and assembly factors prior to export.

**FIG. 1.**
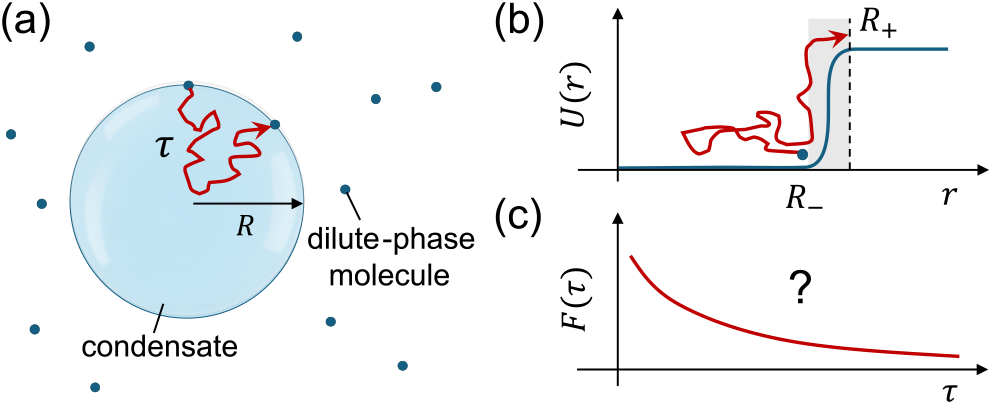
Conceptual picture of single-molecule dwell times in biomolecular condensates. (a) Schematic of a single-molecule tracking (SMT) measurement: a dilute-phase tracer molecule enters a condensate of radius *R* and undergoes stochastic motion before escaping. The dwell time *τ* is the time it spends inside from entry to exit (red trajectory). (b) Illustration of the free-energy (potential-of-mean-force) profile *U* (*r*) as a function of radial coordinate *r*. Escape from the condensate involves both dense-phase diffusion and interfacial barrier crossing. (c) The dwell-time distribution *F* (*τ*) serves as a statistical fingerprint of molecular transport in condensates, motivating a predictive theory for the quantitative interpretation of SMT measurements.

Despite advances in experimental measurements, biophysical models for interpreting SMT data have lagged behind. As a result, SMT dwell times are often fit with phenomenological expressions that lack theoretical justifications. One relevant recent development is the derivation of mean first-passage time for single molecules crossing condensate interfaces [25]; however, the theory is formulated in one dimension and therefore does not directly apply to dwell times in three-dimensional condensates. More importantly, a predictive theory for the full dwell-time distribution—and how its shape maps onto condensate properties such as diffusion coefficients, partitioning, and interfacial kinetics—is still missing [Fig. 1(c)].

Here, we combine SMT experiments on RNA condensates with analytical theory and numerical simulations to identify the physical determinants of molecular dwell times in condensates. We show that dwell-time distributions generically exhibit a power-law regime followed by an exponential decay tail, and that both the power-law exponent and the long-time decay rate depend on whether molecular escape is limited by dense-phase diffusion or interfacial barrier crossing. Our work establishes a quantitative framework for interpreting SMT dwell-time measurements, with implications for understanding molecular transport and retention in both natural and engineered condensates.

## II. RESULTS

### A. Single-molecule tracking of RNA dwell times

To motivate the theoretical analysis, we developed an SMT assay to measure the dwell-time distribution of single RNA transcripts as they enter and exit condensates (Fig. 2). Briefly, RNA molecules transcribed from the *Huntingtin* gene exon1 containing 40 CAG repeats [26] (hereafter f*HTT* −40 RNA) were incubated at a concentration of 3 µM in a phase-separation-promoting buffer [27] to form liquid condensates. Mixing unlabeled f*HTT* −40 RNA with A488-labeled f*HTT* −40 RNA enabled visualization of settled condensates under total internal reflection fluorescence (TIRF) microscopy [Fig. 2(a)]. After condensate sizes stabilized, 5 pM Cy5-labeled f*HTT* −40 RNA was flowed into the microscope chamber for single-molecule tracking [Fig. 2(a)]. Interactions of single Cy5-f*HTT* −40 RNAs with individual condensates were detected as transient fluorescence intensity bursts within the condensate region, from which the dwell time *τ* — the time a molecule remains inside a condensate between entry and exit—was extracted [Fig. 2(b)]. Analysis of the resulting dwell times showed that the distribution *F* (*τ*) follows an approximate power-law decay with an exponent close to − 1.5 over the experimentally accessible window of 0.1–400 s [Fig. 2(c)]. Additional experimental details are provided in the Supplemental Material [28].

**FIG. 2.**
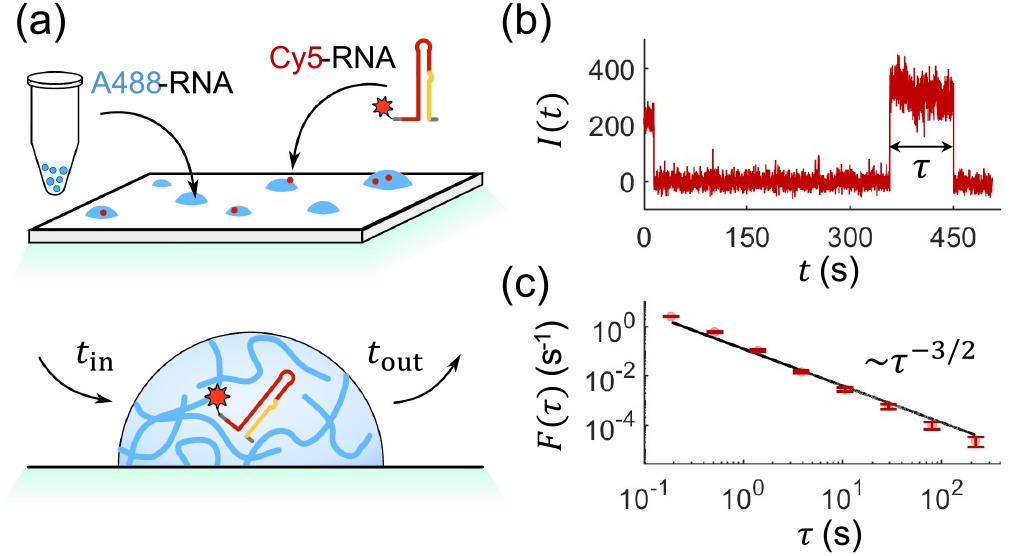
Measuring single-molecule dwell times in phase-separated RNA condensates. (a) Schematic of the SMT assay under TIRF microscopy: A488-labeled RNA condensates were incubated and visualized on a slide before Cy5-labeled RNAs were flowed into the dilute phase (*top*), enabling SMT of the entry and exit times, *t*_in_ and *t*_out_ (*bottom*). (b) Representative time trace following the start of Cy5-RNA flow. A transient intensity burst indicates colocalization of a Cy5-labeled RNA with an A488-labeled condensate. The dwell time is defined as *τ* = *t*_out_*− t*_in_. (c) Measured dwell-time distribution *F* (*τ*) (red circles), with error bars denoting standard deviations estimated by bootstrapping. The black guideline corresponds to a *τ* ^−3*/*2^ power law with normalization prefactor 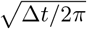, where Δ*t* = 0.1 s is the experimental time resolution (frame interval).

### B. Theoretical formulation of dwell-time distribution

Motivated by the power-law behavior observed in Fig. 2(c), we formulated the dwell time as a first-passage problem: once a molecule enters a condensate, the dwell time is the time it takes to reach the condensate boundary for the first time and exit. To construct a minimal theoretical description, we consider a spherical condensate of radius *R* centered at *r* = 0 in equilibrium with a surrounding dilute phase. By spherical symmetry, the problem depends only on the radial coordinate *r*. Let *p*(*r, t*) denote the probability density that the tracked molecule is at position *r* and time *t* after entry. The time evolution of *p*(*r, t*) is governed by the Smoluchowski equation [25, 29]:

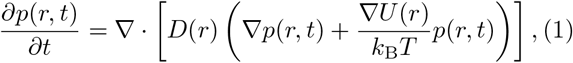

where *D*(*r*) is the position-dependent diffusion coefficient, *U* (*r*) is the potential of mean force experienced by the molecule, *k*_B_ is the Boltzmann constant, and *T* is the temperature. *U* (*r*) is related to the equilibrium concentration profile *c*^eq^(*r*) via

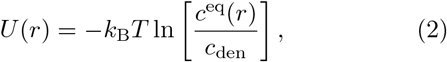

where *c*_den_ is the dense-phase concentration.

The molecule initially enters the condensate from the dilute phase on the spherical shell at *r* = *R*_−_, which translates to the initial condition

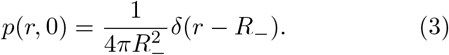

The molecule exits the condensate when it reaches the spherical shell at *r* = *R*_+_, which translates to the absorbing boundary condition

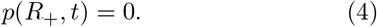

As a natural choice, we set *R*_−_ right inside and *R*_+_ right outside the condensate interface [Fig. 1(b)]. In addition, we have the boundary condition at the condensate center

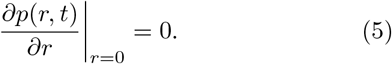

After obtaining *p*(*r, t*) from Eqs. (1)–(5), we compute the survival probability *S*(*t*) by integrating *p*(*r, t*) over the entire condensate. Here, *S*(*t*) is the probability that a molecule remains within the condensate up to time *t*:

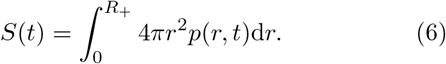

The dwell-time (first-passage-time) distribution is then

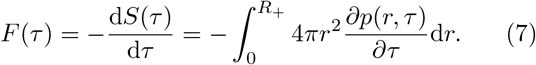

### C. The narrow-interface approximation

Biomolecular condensates typically possess interfaces much narrower than their overall size. Reported interfacial widths are on the order of a few to tens of nanometers [30, 31], whereas condensate radii commonly span hundreds of nanometers to microns. In the narrow-interface limit, spatial variations in the potential of mean force *U* (*r*) and the diffusion profile *D*(*r*) are strongly localized within a thin boundary layer, while the dense interior is essentially homogeneous. This separation of length scales allows us to replace the full Smoluchowski dynamics by a bulk diffusion equation inside the condensate

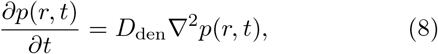

while coarse-graining the transport across the interfacial layer (from *r* = *R*_−_ to *R*_+_) into an effective leaky (Robin) boundary condition:

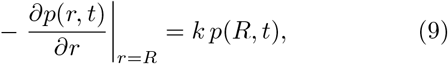

which replaces the absorbing boundary condition in Eq. (4). Here, *k* is an effective permeability parameter: each time a molecule reaches the interface from the dense side, it has a finite probability controlled by *k* to cross the interface. The value of *k* is determined by the shape and height of the barrier as well as the mobility within the interfacial layer:

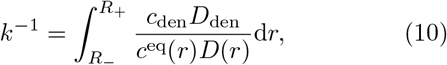

see Supplemental Material for a derivation [28]. In the limit *k*→ +∞, barrier crossing is effectively instantaneous and the boundary becomes absorbing. In the opposite limit *k* →0, the boundary becomes nearly reflective, strongly suppressing escape across the interface.

With the bulk dynamics given by Eq. (8) and the initial and boundary conditions in Eqs. (3), (5), and (9), we derive an analytical solution for *p*(*r, t*):

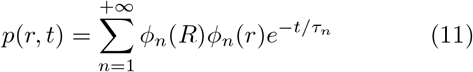

with eigenfunctions

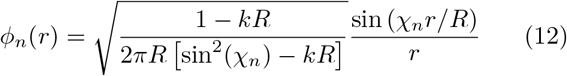

and corresponding decay times

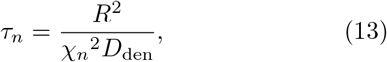

where { *χ*_*n*_} are the positive real roots of the transcendental equation

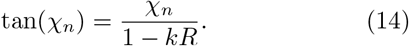

Finally, we obtain *F* (*τ*) in the narrow-interface description by substituting the solution of *p*(*r, t*) into Eq. (7),

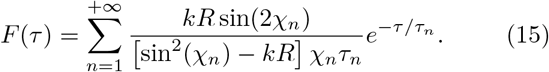

In what follows, we use this framework to discuss the signatures of *F* (*τ*) in the fast and slow barrier-crossing regimes.

### D. Signatures of *F* (*τ*) in the fast interfacial barrier-crossing regime

The presence of an interfacial barrier hinders the escape of molecules once they enter the condensate. The extent of this hindrance depends on how the barrier-crossing timescale compares with the diffusion timescale in the bulk dense phase. When barrier crossing is much faster than dense-phase diffusion, the interface does not appreciably impede molecular escape, corresponding to a large permeability parameter. By taking *kR* ≫1, we Taylor-expand Eq. (14) to leading order in 1*/*(*kR*) and obtain *χ*_*n*_ = *nπ* − *nπ/*(*kR*). Thus

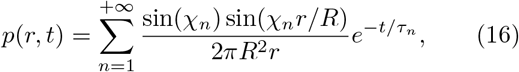

where *τ*_*n*_≈*R*^2^*/*(*n*^2^*π*^2^*D*_den_), and the dwell-time distribution becomes

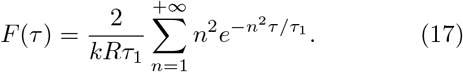

Of particular interest is the relaxation time of the slowest mode *τ*_1_. We refer to this timescale as the dense-phase diffusion timescale *τ*_D_ since it corresponds to diffusive relaxation across the entire condensate:

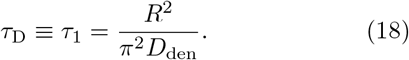

As higher-order modes decay faster (*τ*_*n*_ = *τ*_1_*/n*^2^), *τ*_D_ naturally separates *F* (*τ*) into two asymptotic regimes. (i) For short dwell times (*τ*≪*τ*_D_), the sum in Eq. (17) can be approximated by an integral over *n*, yielding

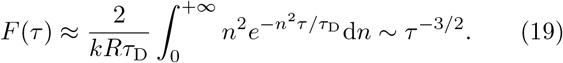

That is, *F* (*τ*) exhibits a 3*/*2 power law in the short-time window. This is consistent with the power-law decay observed in the RNA condensate experiment [Fig. 2(c)]. Physically, this −3*/*2 power law is a universal behavior of 1D diffusive escape near a highly permeable boundary [32, 33], see [28] for a derivation. It applies here because, at short times, molecules remain close to the interface, where the curved boundary is locally flat. (ii) For long dwell times (*τ*≫*τ*_D_), all higher-order modes (*n >* 1) have decayed and the distribution is dominated by the slowest mode, yielding a single-exponential tail

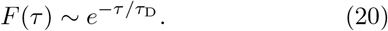

This exponential tail arises from the finite size of the condensate, which sets the timescale *τ*_D_ for molecules to diffuse out. We note that most molecules escape during the −3*/*2 power-law period, since only a small fraction 2*/*(*kR*) ≪ 1 escape via the slowest mode.

### E. Signatures of *F* (*τ*) in the slow interfacial barrier-crossing regime

In the opposite regime, when barrier crossing is much slower than dense-phase diffusion, the interface strongly suppresses molecular escape, corresponding to a small permeability parameter. By taking *kR*≪ 1, we Taylor-expand Eq. (14) and obtain the slowest mode to leading order in 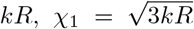. The associated eigen-function is nearly uniform in the condensate interior, 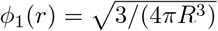. Thus

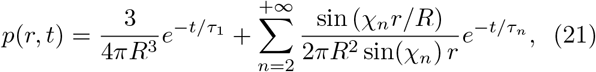

where *τ*_1_ = *R/*(3*kD*_den_), and *χ*_*n*_ ≈ (*n* − 0.5)*π* and *τ*_*n*_ ≈*τ*_D_*/*(*n* −0.5)^2^ for *n* ≥ 2 (see Supplemental Material for a numerical justification [28]).

Physically, the above expression of *p*(*r, t*) reflects two processes with a clear separation of timescales. The higher-mode sum captures rapid diffusion-driven relaxation that smooths out spatial gradients inside the condensate over a timescale *τ*_D_, after which the profile is essentially uniform. The leading (uniform) mode then decays much more slowly due to rare escape across the barely permeable interface. We denote this slow timescale by *τ*_B_ since it is controlled by interfacial barrier crossing:

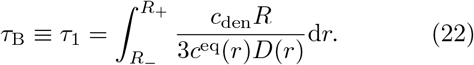

Note that the slow barrier-crossing regime implies *kR* ~ *τ*_D_*/τ*_B_ ≪ 1.

The corresponding dwell-time distribution becomes

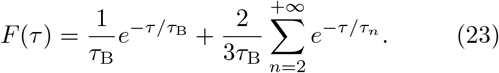

As discussed above, *τ*_D_ (more precisely *τ*_2_ ≈ *τ*_D_*/*2) naturally separates *F* (*τ*) into two asymptotic regimes. (i) For short dwell times (*τ*≪*τ*_D_), the sum in Eq. (23) can be approximated by an integral over *n*, yielding

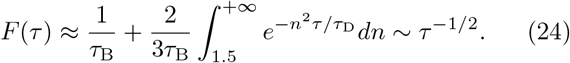

That is, *F* (*τ*) exhibits a − 1*/*2 power law in the short-time window, which is expected for 1D diffusive escape near a barely permeable boundary, as derived in the Supplemental Material [28]. (ii) For long dwell times (*τ*≫*τ*_D_), all higher-order modes (*n* ≥ 2) have decayed, yielding a single-exponential tail

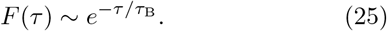

This exponential tail reflects rare interfacial escape after molecules have become well mixed within the condensate. Here, most molecules escape during the exponential decay period, since the fraction of molecules that escape through the slowest mode is approximately 1.

### F. Numerical validation

To test our theoretical predictions for biological condensates with finite-width interfaces, we specified radial profiles for the equilibrium concentration and diffusivity with a smooth transition across the interface [30]:

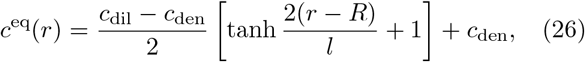

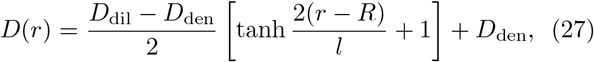

where *c*_den*/*dil_ and *D*_den*/*dil_ are the dense-/dilute-phase concentration and diffusion coefficient, respectively, and *l* is the interface width. We then numerically solved the full Smoluchowski framework, Eqs. (1)–(5), and computed *F* (*τ*) using Eq. (7).

In the fast barrier-crossing regime (parameter set 1 in Table I), we show *F* (*τ*) for a droplet of radius *R* = 1 µm in Fig. 3(a) (blue). As expected, *F* (*τ*) follows a −1.5 power law at short times before crossing over to a single-exponential tail, with the transition occurring around *τ*_D_. We further solved for *F* (*τ*) at several values of *R* and extracted the long-term decay time from the exponential tail for each *R*. These decay times agree with the predicted timescale *τ*_D_ in Eq. (18) and scale as *R*^2^ across condensate sizes [Fig. 3(b)], suggesting that the dwell time is controlled by dense-phase diffusion. Since most molecules escape during the − 1.5 power-law regime, their dwell times are typically smaller than *τ*_D_. Thus, molecules often leave the condensate before fully exploring the droplet interior, as illustrated by the representative trajectories in Fig. 3(c).

**TABLE I.**
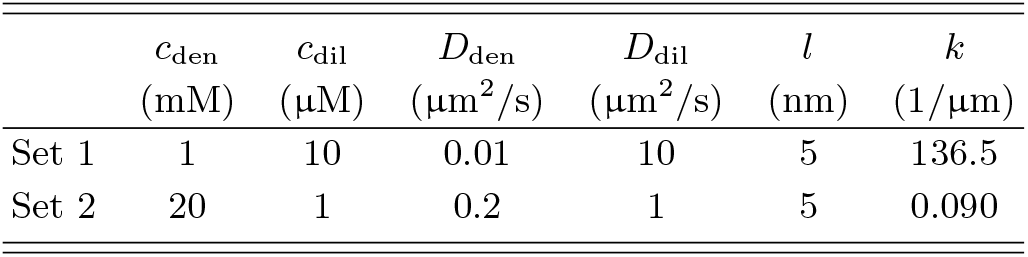
Parameters for numerical simulations.

**FIG. 3.**
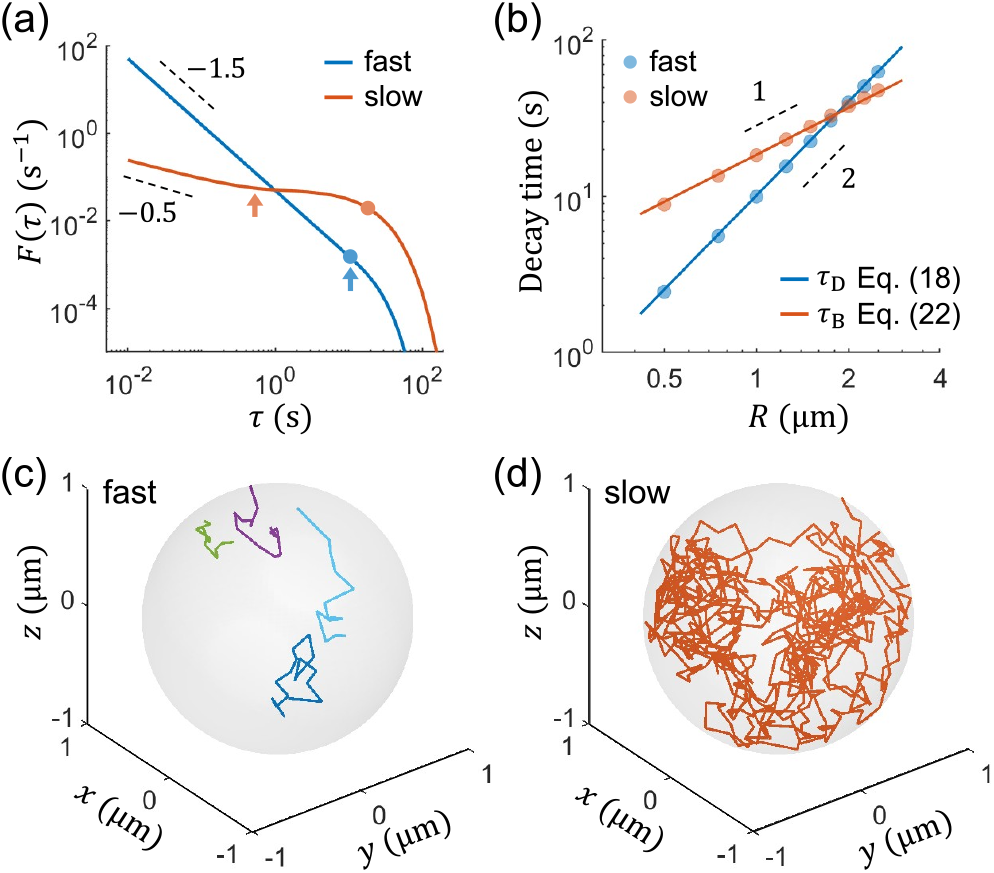
Numerical demonstration of signatures for fast and slow interfacial barrier crossing. (a) Numerical solutions for *F* (*τ*) in the fast (blue) and slow (orange) barrier-crossing regimes for a droplet of radius *R* = 1 µm. Arrows indicate the crossover time *τ*_D_, at which the distribution transitions from a short-time power law to a long-time exponential decay. Circles denote the decay times of the exponential tails obtained by numerical fitting. (b) Decay time versus droplet radius *R*. Circles are numerical results, and curves are theoretical predictions. (c) and (d) Representative trajectories for *R* = 1 µm in the fast and slow barrier-crossing regimes, respectively. Each color denotes a different trajectory. Trajectories were obtained by simulating the overdamped Langevin equation corresponding to Eq. (1) with parameters listed in Table I. For all panels, we used *R*_*±*_ = *R ±* 3*l*. Additional numerical details are provided in the Supplemental Material [28].

In contrast, in the slow barrier-crossing regime (parameter set 2 in Table I), *F* (*τ*) for *R* = 1 µm exhibits a − 0.5 power law followed by a single-exponential decay, Fig. 3(a) (orange). Again, the transition occurs around *τ*_D_, but the exponential tail decays on a much longer timescale. The extracted long-term decay time versus *R* agrees with the predicted barrier-crossing timescale *τ*_B_ in Eq. (22) and grows linearly with *R* [Fig. 3(b)], suggesting that the dwell time is dominated by interfacial barrier crossing. Since most molecules escape during the exponential-decay regime, their dwell times are typically on the order of *τ*_B_≫*τ*_D_. Thus, molecules often explore the entire droplet multiple times before escaping, as illustrated by the representative trajectory in Fig. 3(d).

### G. Interpreting RNA dwell times

Having established the theoretical signatures of the two escape regimes, we now use them to interpret the measured RNA dwell-time distribution in Fig. 2(c). The approximate − 1.5 power law observed in the RNA dwell-time distribution is consistent with fast interfacial-barrier crossing, with escape primarily controlled by dense-phase diffusion in the measured RNA condensates. In this regime, our theory predicts that *F* (*τ*) should eventually cross over to an exponential tail on a timescale *τ*_D_ = *R*^2^*/*(*π*^2^*D*_den_). While direct resolution of this tail is limited by Cy5 photobleaching in the present assay (lifetime ~ 210 s), the absence of a clear exponential tail in the current time window suggests *τ*_D_ *>* 100 s, corresponding to *D*_den_ *<* 2.5 × 10^−4^ µm^2^*/*s for micron-sized droplets in our experiments (*R* ≈ 0.5 µm). This small upper bound on *D*_den_ agrees with prior studies showing that RNA-rich condensates, particularly those formed by repeat-containing RNAs, can exhibit strongly restricted molecular mobility [27, 34], supporting the interpretation that RNA escape in the present condensates is dense-phase diffusion limited.

## III. DISCUSSION

Motivated by the experimental measurements of single-RNA dwell times in condensates, we developed a theoretical framework for molecular residence times. We show that the dwell-time distribution *F* (*τ*) generically exhibits a short-time power-law regime followed by a long-time exponential decay. The form of this distribution reflects the rate-limiting mechanism of molecular escape: diffusion-limited dynamics yield a − 1.5 power law with an exponential tail set by the diffusion timescale *τ*_D_ ~ *R*^2^, whereas interface-limited dynamics yield a − 0.5 power law with a decay governed by the barrier-crossing timescale *τ*_B_ ~ *R*. These distinct signatures provide a direct and experimentally accessible means to infer the dominant transport mechanism in condensates from single-molecule measurements.

Our results suggest that biological condensates can operate in different regimes depending on their physical properties. A useful dimensionless parameter is *kR* ~ *τ*_D_*/τ*_B_, which compares the timescale of dense-phase diffusion to that of interfacial escape. Droplets with low interfacial permeability are more likely to fall into the slow barrier-crossing regime, where molecules “equilibrate” within the condensate before escaping. In contrast, in higher-permeability droplets, molecules often escape before sampling the full condensate interior. This distinction may have important biological consequences: molecules that fully explore the condensate can sample many potential binding partners or reaction sites, thereby enhancing reaction efficiency, whereas those that only partially explore it may be limited to local or surface-proximal interactions.

Our theoretical framework is intentionally minimal, allowing us to obtain analytical predictions for dwell-time distributions while isolating the roles of dense-phase diffusion and interfacial escape. Biomolecular condensates, especially in cellular contexts, can exhibit additional complexity, including heterogeneous organization, network formation, viscoelastic relaxation, aging, and active remodeling [18, 20, 24, 35, 36]. We hope that this work will motivate future studies to incorporate these effects and establish dwell-time distributions as quantitative probes of the physical mechanisms governing molecular transport and retention in biomolecular condensates.

## DATA AVAILABILITY

The data supporting the findings of this study, including the experimental dwell-time measurements, numerical results, and analysis code used to generate the figures, are available in [repository name] at DOI: [DOI].

## ACKNOWLEDGMENTS

We thank Robert L. Leheny, Daniel H. Reich, Armaan Ahmed, and Vladimir Grigorev for insightful discussions and valuable feedback on the manuscript. F.Y., C.W., and Y.Z. were supported by a startup fund at Johns Hopkins University and by NIH Award No. R35GM162296.

Y.Z. acknowledges support from the Sloan Foundation through a Sloan Research Fellowship (FG-2025-25076). R.M., M.L.R., and S.A.W. were supported by NIH Award Nos. R35GM136351 and R21NS128701. This work was carried out at the Advanced Research Computing at Hopkins (ARCH) core facility (rockfish.jhu.edu), which is supported by NSF Grant No. OAC1920103.

## Notes

### Competing Interest Statement

The authors have declared no competing interest.

