## Supplemental for "Single-Molecule Dwell Times in Biomolecular Condensates"

(Dated: June 29, 2026)

### I. EXPERIMENTAL METHODS

#### A. RNA preparation

f*HTT*40 RNA containing the repeat region of human *HTT* exon 1 with 40 CAG triplets and flanking residues [1] was transcribed *in vitro* with T7 RNAP and 2 mM ATP, 2 mM UTP, 6 mM GTP, 6 mM CTP. The RNA was purified by denaturing 5% polyacrylamide PAGE and stored in water at −80 °C for 1 month. For random labeling with ATTO-488, RNA was transcribed as described above except using 2 mM ATP, 0.5 mM UTP, 6 mM GTP, 6 mM CTP and 10–40 µM fluorophore-5-aminoallyl-UTP, then conjugated with ATTO-488 NHS-ester.

5′-Cy5-labeled f*HTT*40 RNA was prepared by transcribing RNA with 1 mM ATP, 1 mM UTP, 5 mM GTP, 5 mM CTP, 30 mM GMP (250 µL total volume), at 37 °C for 2 h. The RNA was purified by denaturing PAGE as above and resuspended in nuclease-free water to <15 µM (final). Previously established protocols [2, 3] were followed to generate a 5′ primary amine, in which 37.5 µL 15 µM 5′-GMP labeled RNA was mixed with 37.5 µL 2X buffer (10 mM potassium phosphate, pH 7, 150 mM NaCl, 10 mM EDTA final). To this was added 12.5 mg 1-ethyl-3-(3-dimethylaminopropyl) carbodiimide (EDC), 50 µL 0.25 M ethylenediamine dihydrochloride and 200 µL 0.1 M imidazole at pH 6. The reaction was incubated at 37 °C for 3 h. The RNA was precipitated with ethanol and resuspended in 20 µL 0.1 M sodium carbonate buffer, pH 8.5–9. To this was added 30 µL Cy5-monoreactive dye (Cytiva) overnight at room temperature. The excess unreacted dye was removed using a size-exclusion column (Takara) followed by ethanol precipitation and resuspension in nuclease free water. The RNA was stored at −80 °C for up to one month. The percentage of RNA labeled with Cy5 was determined by absorption using $\varepsilon_{650}$ (Cy5) = 250,000 $M^{-1}cm^{-1}$ and $\varepsilon_{260}$ for each RNA after correction for A260 of Cy5 (0.05). A small fraction of the RNA was repurified (Zymo RNA cleanup kit) and the percentage of labeled RNA was confirmed to be >95%.

#### B. Single RNA binding assays

***TIRF microscopy.*** Association of single f*HTT*40 RNAs with RNA droplets was monitored on a custom-built prism total internal reflection fluorescence (TIRF) microscope with an EMCCD camera (Andor) [4]. Video (100 ms/frame) was acquired with 488 nm and 640 nm excitation of ATTO-488 and Cy5 fluorophores, respectively, using custom scripts implemented in IDL (smCamera, https://github.com/Ha-SingleMoleculeLab/Data-Aquisition). Samples were deposited on quartz slides passivated using dodecylsilane and Tween-20 [5]. All imaging was performed at 21 °C.

***Phase separation and droplet imaging.*** To induce liquid-liquid phase separation, f*HTT*40 RNA (50 µL 3 µM solution in water) was heated to 65 °C for 1 min and refolded on ice for 1 min and at room temperature for 1 min. The refolded RNA was mixed with an equal volume of buffer to a final concentration of 10 mM Tris-HCl pH 7.5, 1 mM $MgCl_2$, 750 mM NaCl, and 15% PEG 8000 (1X phase separation buffer) [6]. 1 µL RNase inhibitor, 2 µL glucose, and 4 µL glucose oxidase (oxygen scavenging system, OSS) were added to the 100 µL droplet mixture, and the mixture was immediately added to a channel on the passivated slide. After the droplets were allowed to settle for 10 min, 100 µL 10–20 nM ATTO-488-f*HTT*40 RNA refolded and prepared as above was added to the channel. This process was repeated once. The channel was then washed twice with 100 µL 1X phase separation buffer containing 15% PEG 8000 with OSS. The ATTO-488-f*HTT*40 RNA was incorporated into the unlabeled droplets, allowing settled droplets to be detected upon excitation at 488 nm. Droplet sizes were determined using 3D Objects Counter in ImageJ and a pixel size of 0.15 µm [7]. The droplet sizes at 15 min and 1 h remained constant, indicating that the droplets were stable under these conditions.

***RNA-droplet binding kinetics.*** To observe the association of single RNAs with settled droplets, 5 pM refolded 5′-Cy5-f*HTT*40 RNA in 1X phase separation buffer containing 15% PEG 8000 with OSS was added to the slide channel with settled droplets immediately after beginning 640 nm excitation. The start of flow was

---

* Present address: The Laboratory of Biochemistry and Genetics, The National Institute of Diabetes and Digestive and Kidney Diseases, The National Institutes of Health, Bethesda, Maryland 20892, USA
† Contact author:

marked as the beginning of the experiment. The droplets were imaged again (∼2–10 s) with 488 nm excitation to determine their positions after the flow of 5′-Cy5-RNA solution. Excitation was then switched to 640 nm (Cy5) for 10 min (10 frames/s). The dead time between the addition of 5′-Cy5-RNA and the start of Cy5 excitation was 2–15 s for all experiments. At the end of the movie, the droplets were again imaged with the 488 nm laser for ∼2 s.

***Single molecule data analysis.*** Movies were analyzed using Imscroll [8]. Areas of interest (AOIs) in each field of view, corresponding to the areas occupied by individual droplets, were defined by the fluorescence intensity upon 488 nm excitation. For each droplet (AOI), 5′-Cy5-f*HTT*40 RNA binding events were obtained from changes in the time-resolved Cy5 fluorescence intensity as described previously [8]. Fits of individual trajectories yielded the times of association ($t_{\rm in}$), dissociation ($t_{\rm out}$), and the residence lifetimes ($\tau$) of each binding event. For simplicity, only single, non-overlapping binding events were included in the analysis. The contribution of photobleaching to the measured residence times was estimated from the loss of fluorescence of immobilized 5′-Cy5-f*HTT*40 RNAs over time under the identical conditions, yielding $\tau_{\rm PB} = 210$ s. We confirmed that free Cy5 dye does not appreciably interact with the droplets under these conditions.

### II. THEORETICAL METHODS

#### A. Derivation of the effective interfacial permeability $k$ in Eq. (10)

The derivation of $k$ follows closely the approach developed in [9]. In the narrow-interface regime, we can assume that the flux through the interface is uniform in space, i.e.,

$$j(t) = -D(r)\left[\frac{\partial p(r,t)}{\partial r} + \frac{1}{k_{\rm B}T}\frac{{\rm d}U(r)}{{\rm d}r}p(r,t)\right]. \tag{S1}$$

Thus,

$$\frac{\partial p(r,t)}{\partial r} + \frac{1}{k_{\rm B}T}\frac{{\rm d}U(r)}{{\rm d}r}p(r,t) = -\frac{j(t)}{D(r)}. \tag{S2}$$

The solution to the above equation is

$$\begin{aligned} p(r,t) &= -j(t)\int_{R_+}^{r} \exp\left[\frac{U(r') - U(r)}{k_{\rm B}T}\right]\frac{{\rm d}r'}{D(r')} \\ &= j(t)\int_{r}^{R_+} \frac{c^{\rm eq}(r)\,{\rm d}r'}{c^{\rm eq}(r')D(r')}. \end{aligned} \tag{S3}$$

We assume that the interface spans $r \in [R_-, R_+]$. It then follows that $c_{\rm den} = c^{\rm eq}(R_-)$ and $D_{\rm den} = D(R_-)$. The flux at $r = R_-$ is continuous, which gives

$$j(t) = -D_{\rm den}\left.\frac{\partial p(r,t)}{\partial r}\right|_{r=R_-} = D_{\rm den}kp(R_-,t), \tag{S4}$$

where the first and second terms are the fluxes from the interfacial and dense-phase sides, respectively, and the last term comes from the Robin boundary condition in Eq. (9). Plugging Eq. (S3) into Eq. (S4) gives

$$j(t) = D_{\rm den}kj(t)\int_{R_-}^{R_+} \frac{c_{\rm den}\,{\rm d}r}{c^{\rm eq}(r)D(r)}. \tag{S5}$$

From which we obtain an expression for the interface permeability $k$,

$$k^{-1} = \int_{R_-}^{R_+} \frac{c_{\rm den}D_{\rm den}}{c^{\rm eq}(r)D(r)}{\rm d}r. \tag{S6}$$

#### B. Origins of the short-time power laws

At short times, molecules sample only regions near the condensate interface, so the curved boundary can be treated as locally flat. We therefore consider 1D diffusion on the half-line $x > 0$, where $x$ is the distance from the interface into the condensate.

First, consider the high-permeability limit, in which the interface is effectively absorbing. For a molecule initially located at $x = x_0$, the probability density satisfies

$$\frac{\partial p}{\partial t} = D_{\rm den}\frac{\partial^2 p}{\partial x^2}, \tag{S7}$$

with $p(x,0) = \delta(x - x_0)$ and $p(0,t) = 0$. Using the method of images, the absorbing-boundary solution is

$$p(x,t) = \frac{1}{\sqrt{4\pi D_{\rm den}t}}\left[e^{-\frac{(x-x_0)^2}{4D_{\rm den}t}} - e^{-\frac{(x+x_0)^2}{4D_{\rm den}t}}\right]. \tag{S8}$$

The escape-time (dwell-time) distribution is the outward flux through the interface,

$$F(\tau) = D_{\rm den}\left.\frac{\partial p}{\partial x}\right|_{x=0} = \frac{x_0}{\sqrt{4\pi D_{\rm den}\tau^3}}e^{-\frac{x_0^2}{4D_{\rm den}\tau}}, \tag{S9}$$

which reduces to

$$F(\tau) \sim \tau^{-3/2} \tag{S10}$$

for small $x_0$. Thus, 1D diffusive escape near a highly permeable boundary gives the $-3/2$ power law.

Next, consider the opposite limit of an interface with very low permeability. Escape through the interface is described by the boundary condition

$$\left.\frac{\partial p}{\partial x}\right|_{x=0} = k\,p(0,t). \tag{S11}$$

When $k$ is small, the short-time concentration profile is well approximated by the reflecting-boundary solution. Using the method of images,

$$p(x,t) = \frac{1}{\sqrt{4\pi D_{\rm den}t}}\left[e^{-\frac{(x-x_0)^2}{4D_{\rm den}t}} + e^{-\frac{(x+x_0)^2}{4D_{\rm den}t}}\right]. \tag{S12}$$

The probability density at the interface is therefore

$$p(0,t) = \frac{1}{\sqrt{\pi D_{\text{den}} t}} e^{-\frac{x_0^2}{4D_{\text{den}} t}}, \tag{S13}$$

which is approximately

$$p(0,t) \approx \frac{1}{\sqrt{\pi D_{\text{den}} t}} \tag{S14}$$

for small $x_0$. The escape-time distribution is therefore

$$F(\tau) = D_{\text{den}} \left. \frac{\partial p}{\partial x} \right|_{x=0} = D_{\text{den}} k p(0,\tau) \sim \tau^{-1/2}. \tag{S15}$$

Thus, 1D diffusive escape near a barely permeable boundary gives the $-1/2$ power law.

#### C. Numerical justification for $\chi_n \approx (n-0.5)\pi$ in Eq. (21)

In the slow interfacial barrier-crossing regime ($kR \ll 1$), the transcendental equation in Eq. (14) reduces to

$$\tan(\chi_n) = \chi_n. \tag{S16}$$

For $n \geq 2$, $\chi_n$ becomes sufficiently large that the roots of the above equation lie close to the positive poles of $\tan(\chi_n)$, giving

$$\chi_n \approx (n-0.5)\pi. \tag{S17}$$

In Fig. S1, we show the comparison between the exact and approximated values of $\chi_n$ for $n \geq 2$ at a few choices of $kR$. The case $kR = 0.1$ is close to that of parameter set 2, $kR = 0.09$, used in the main text. The exact solution is obtained numerically as the $n$th positive root of Eq. (14) in the interval $\chi_n \in \left[(n-1)\pi, (n-0.5)\pi - 10^{-6}\right]$, where the small offset of $10^{-6}$ is used to avoid singularities at the poles of $\tan(\chi_n)$. The approximation agrees well with the exact solution.

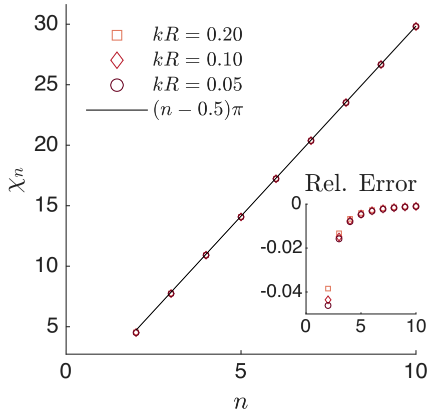


FIG. S1. Numerical validation of $\chi_n \approx (n-0.5)\pi$ in Eq. (21) for $n \geq 2$ at several choices of $kR$ values: 0.05, 0.1, and 0.2. Symbols denote exact numerical solutions of the transcendental equation, Eq. (14), and the black line denotes the analytical approximation. Inset: Relative error of the approximation for the same $kR$ values.

#### D. Numerical details for solving $F(\tau)$

To obtain the numerical solution of $F(\tau)$, we first computed $p(r,t)$ by solving the radially symmetric Smoluchowski equation in Eq. (1) with initial and boundary conditions in Eqs. (3)–(5) and the radial profiles for the equilibrium concentration and diffusivity specified in Eqs. (26) and (27). For a spherical condensate of radius $R$ and interface width $l$, we used a piecewise-defined radial grid: a coarse spacing $\Delta r_{\text{bulk}} = 0.001\,\mu\text{m}$ in the uniform bulk ($r \leq R_- - 2l$), and a fine spacing $\Delta r_{\text{int}} = l/50$ in the transition region ($R_- - 2l < r < R_+$) to improve numerical accuracy. The Smoluchowski equation was solved using MATLAB's *pdepe* function, which employs the above-mentioned spatial discretization and a variable-step, variable-order time integrator [10, 11].

After obtaining $p(r,t)$, we computed the survival probability $S(t)$ by integrating over the condensate, as in Eq. (6), and obtained the dwell-time distribution $F(\tau)$ by numerically differentiating $S(t)$ with $dt = 0.01\,\text{ms}$, as in Eq. (7). For each parameter set and droplet radius, we computed $F(\tau)$ over the time interval $[0.01\,\text{s}, \tau_{\max}]$, where $\tau_{\max} = 20\max(\tau_{\text{D}}, \tau_{\text{B}})$, to ensure that the exponential tail was well captured. We then renormalized $F(\tau)$ so that its integral over the simulated time interval was equal to 1.

To obtain the scaling relationship between the condensate radius $R$ and the decay time, we repeated the above procedure for various $R$ while keeping $l$ fixed and setting $R_\pm = R \pm 3l$. To extract the fitted decay time of the exponential tail, $\tau_{\text{fit}}$, we fitted $F(\tau)$ over the interval $\tau \in [2\max(\tau_{\text{D}}, \tau_{\text{B}}), \tau_{\max}]$ to a single-exponential decay function. The lower bound of the interval was chosen to ensure that all higher order modes of $F(\tau)$ had sufficiently decayed.

#### E. Numerical details for simulating the overdamped Langevin equation

The representative trajectories in Figs. 3(c) and 3(d) were obtained by simulating the overdamped Langevin equation in three dimensions [12],

$$\frac{\text{d}\mathbf{r}}{\text{d}t} = -\frac{D(r)}{k_{\text{B}}T}\nabla U(r) + \nabla D(r) + \boldsymbol{\eta}(r,t), \tag{S18}$$

which provides a single-molecule-level stochastic description of the ensemble-level Smoluchowski equation in Eq. (1). Here, $\boldsymbol{\eta}(r,t)$ denotes Gaussian white noise with independent Cartesian components satisfying

$$\langle \eta_i(t) \rangle = 0, \tag{S19}$$

$$\langle \eta_i(t)\eta_j(t') \rangle = 2D(r)\delta_{ij}\delta(t - t'), \tag{S20}$$

where $i, j \in \{x, y, z\}$. Physically, the first term on the right-hand side of Eq. (S18) describes deterministic drift due to the potential of mean force, the second term is the spurious drift required for the position-dependent diffusivity to ensure detailed balance, and the last term describes thermal fluctuations.

We simulated single-particle trajectories using radial profiles $U(r)$ and $D(r)$ specified by Eqs. (2), (26), and (27), with parameters taken from Table 1 for a condensate of radius $R = 1\,\mu\text{m}$. Each particle starts at $r = R_-$, and its position is updated according to

$$\mathbf{r}(t + \Delta t) = \mathbf{r}(t) - \frac{D(r)}{k_\mathrm{B}T}\nabla U(r)\,\Delta t + \nabla D(r)\,\Delta t + \sqrt{2D(r)\Delta t}\,\boldsymbol{g}, \tag{S21}$$

where the components of $\boldsymbol{g}$ are independent Gaussian random variables with mean 0 and variance 1. The simulation timestep was $\Delta t = 0.01\,\mu\text{s}$ and particle positions were recorded every 0.1 ms. Each trajectory ends when the particle reaches $r = R_+$.

[10] The MathWorks, Inc., MATLAB, version r2025b, Natick, Massachusetts, USA (2025).

[11] The MathWorks, Inc., pdepe: Solve initial-boundary value problems for parabolic–elliptic pdes in 1-d, MATLAB Documentation (2026).

[12] S. Bo, L. Hubatsch, J. Bauermann, C. A. Weber, and F. Jülicher, Stochastic dynamics of single molecules across phase boundaries, Physical Review Research **3**, 043150 (2021).
